# Restoring AIBP expression in the retina provides neuroprotection in glaucoma

**DOI:** 10.1101/2023.10.16.562633

**Authors:** Won-Kyu Ju, Yonju Ha, Seunghwan Choi, Keun-Young Kim, Tonking Bastola, Jungsu Kim, Robert N. Weinreb, Wenbo Zhang, Yury I. Miller, Soo-Ho Choi

**Affiliations:** Hamilton Glaucoma Center and Shiley Eye Institute, The Viterbi Family Department of Ophthalmology, University of California San Diego, La Jolla, CA 92039, USA; Department of Ophthalmology and Visual Sciences, University of Texas Medical Branch, TX, 77555-0144, USA; National Center for Microscopy and Imaging Research, Department of Neurosciences, University of California San Diego, La Jolla, CA 92039, USA; Department of Medicine, University of California San Diego, La Jolla, CA 92039, USA; Raft Pharmaceuticals LLC, San Diego, CA 92121, USA

**Keywords:** Glaucoma, AIBP, AAV, RGC protection, TLR4-lipid raft

## Abstract

Glaucoma is a neurodegenerative disease manifested in retinal ganglion cell (RGC) death and irreversible blindness. While lowering intraocular pressure (IOP) is the only proven therapeutic strategy in glaucoma, it is insufficient for preventing disease progression, thus justifying the recent focus on targeting retinal neuroinflammation and preserving RGCs. We have identified apolipoprotein A-I binding protein (AIBP) as the protein regulating several mechanisms of retinal neurodegeneration. AIBP controls excessive cholesterol accumulation via upregulating the cholesterol transporter ATP-binding cassette transporter 1 (ABCA1) and reduces inflammatory signaling via toll-like receptor 4 (TLR4) and mitochondrial dysfunction. ABCA1, TLR4 and oxidative phosphorylation components are genetically linked to primary open-angle glaucoma. Here we demonstrated that AIBP and ABCA1 expression was decreased, while TLR4, interleukin 1 β (IL-1β), and the cholesterol content increased in the retina of patients with glaucoma and in mouse models of glaucoma. Restoring AIBP expression by a single intravitreal injection of adeno-associated virus (AAV)-AIBP protected RGCs in glaucomatous DBA/2J mice, in mice with microbead-induced chronic IOP elevation, and optic nerve crush. In addition, AIBP expression attenuated TLR4 and IL-1β expression, localization of TLR4 to lipid rafts, reduced cholesterol accumulation, and ameliorated visual dysfunction. These studies collectively indicate that restoring AIBP expression in the glaucomatous retina reduces neuroinflammation and protects RGCs and Müller glia, suggesting the therapeutic potential of AAV-AIBP in human glaucoma.

**Significance Statement:** Lowering intraocular pressure is insufficient to prevent progressive neuroinflammation, retinal ganglion cell (RGC) death and vision loss in glaucoma. We show that reduced expression of AIBP is associated with increased markers of neuroinflammation, TLR4 and IL-1β, in human glaucomatous retina and mouse models of glaucoma. Restoring AIBP expression by an intravitreal injection of AAV-AIBP protects RGCs and preserves visual function in mouse models of glaucoma. Mechanistically, we demonstrate that AAV-AIBP disrupts TLR4-lipid raft formation and attenuates phosphorylation of AMPK, ERK1/2 and p38 and expression of TLR4 and IL-1β *in vivo*. Collectively, our findings suggest that AIBP-mediated disruption of retinal TLR4-lipid rafts can prevent RGC loss and demonstrate the therapeutic potential of AAV-AIBP in treatment of human glaucoma.

## Introduction

Glaucoma is a leading cause of blindness worldwide and is characterized by a slow, progressive, and irreversible degeneration of retinal ganglion cells (RGCs) and RGC axons, leading to vision loss (1). Pathophysiological processes leading to the development of glaucoma include elevated intraocular pressure (IOP), glia-driven neuroinflammation, mitochondrial dysfunction, oxidative stress, and vascular dysfunction (2). Currently, elevated IOP is the only proven modifiable risk factor; however, lowering IOP is often not sufficient to prevent glaucoma progression.

Activation of toll-like receptor-4 (TLR4) has been reported as a major component of retinal neuroinflammation. TLR4 polymorphism is associated with the higher risk of glaucoma (3). Elevated IOP increases TLR4 expression, activates inflammasome-associated signaling and induces interleukin-1β (IL-1β)-mediated inflammatory response in retinal degeneration (4, 5). In addition, TLR4 activation is associated with mitochondrial damage caused by intracellular reactive oxygen species (ROS) and defective mitochondrial dynamics (6). This agrees with the report of a polymorphism in mitochondrial cytochrome c oxidase subunit I of the oxidative phosphorylation (OXPHOS) complex (Cx)-IV associated with the development of glaucoma (7). In addition, mitochondrial dysfunction and metabolic stress induced by glaucomatous insults such as elevated IOP result in RGC degeneration (2, 8–10). Genetic deletion or pharmacological inhibition of TLR4 significantly reduces RGC death and proinflammatory responses in experimental glaucoma (11, 12). Our recent study demonstrated that TLR4 and IL-1β expression is upregulated in Müller glia in glaucomatous human and DBA/2J mouse retinas (13). However, clinical development of specific antagonists of TLR4 for multiple indications was so far unsuccessful.

TLR4 activation requires its localization to and homodimerization in lipid rafts, ordered membrane domains with high content of cholesterol and sphingolipids (14–16). Removal of cholesterol from lipid rafts in activated cells disrupts TLR4 signaling (17). Conversely, cholesterol loading of the plasma membrane can activate TLR4-dependent signaling pathways (18). The covariance of membrane cholesterol and TLR4 activation implies that disease-associated dysregulation of cellular cholesterol metabolism and its excessive accumulation may contribute to TLR4 activation. Thus, selective cholesterol depletion from cells expressing high levels of TLR4 would inhibit TLR4 activation and neuroinflammation in glaucoma.

Apolipoprotein A-I binding protein (AIBP; gene name *APOA1BP* or *NAXE*) is a secreted protein that binds apolipoprotein A-I, stabilizes the cholesterol transporter ABCA1 and augments cholesterol efflux (19–21). In addition, AIBP binds to TLR4 and thus directs cholesterol depletion and disruption of lipid rafts to TLR4-expressing cells (14, 22). Recent studies from our group demonstrated that in mouse retina, AIBP was predominantly expressed in RGCs and that AIBP deficiency exacerbated RGC vulnerability to cell death and triggered vision impairment in response to elevated IOP (13). Furthermore, AIBP deficiency was associated with mitochondrial dysfunction in both RGCs and Müller glia and with glia-driven upregulation of the inflammatory TLR4/ IL-1β signaling in glaucomatous neurodegeneration (13).

In this study, we found decreased ABCA1 and AIBP expression that was associated with increased TLR4/IL-1β expression and cholesterol deposition in the retina of patients with glaucoma and in mouse models of glaucoma. Using intravitreal delivery of adeno-associated virus (AAV)-AIBP, we demonstrated that restoring AIBP expression inhibited TLR4 activation and IL-1β expression, reduced retinal cholesterol accumulation, protected RGCs and their axon, and ameliorated visual dysfunction in the glaucomatous mouse retina. These results provide *in vivo* evidence that sustained AIBP expression is a key regulator of neuroprotection and may constitute a novel therapeutic strategy for RGC survival and vision preservation in glaucoma.

## Results

### Reduced expression of ABCA1 and AIBP in human glaucomatous retina

Since ABCA1 has been implicated in glaucoma (25), we conducted an immunohistochemical (IHC) survey of retina specimens from human donors without and with glaucoma diagnosis. The glaucoma diagnosis was confirmed by documenting a significant loss of RGCs and their axons in the optic nerve head (ONH) (*SI Appendix, Fig. S1*). In normal human retina, we observed ABCA1 immunoreactivity in all retinal layers, predominantly localized in RGC somas and axons labeled with neuron-specific β-III tubulin (TUJ1) (Figs. 1 A), and in Müller glia labeled with glutamine synthase (GS) (*SI Appendix,* Fig. S2A). In human glaucomatous retina, the ABCA1 expression was reduced in all retinal layers and significantly diminished in RGCs and Müller glia (Figs. 1 A and C; *SI Appendix,* Fig. S2A). We also found that the patterns of AIBP expression were similar to those for ABCA1 in the control and glaucomatous human retinas and that AIBP was also significantly decreased in the glaucomatous RGCs and Müller glia (Figs. 1 B and C; *SI Appendix,* Fig. S2B).

**Fig. 1.**
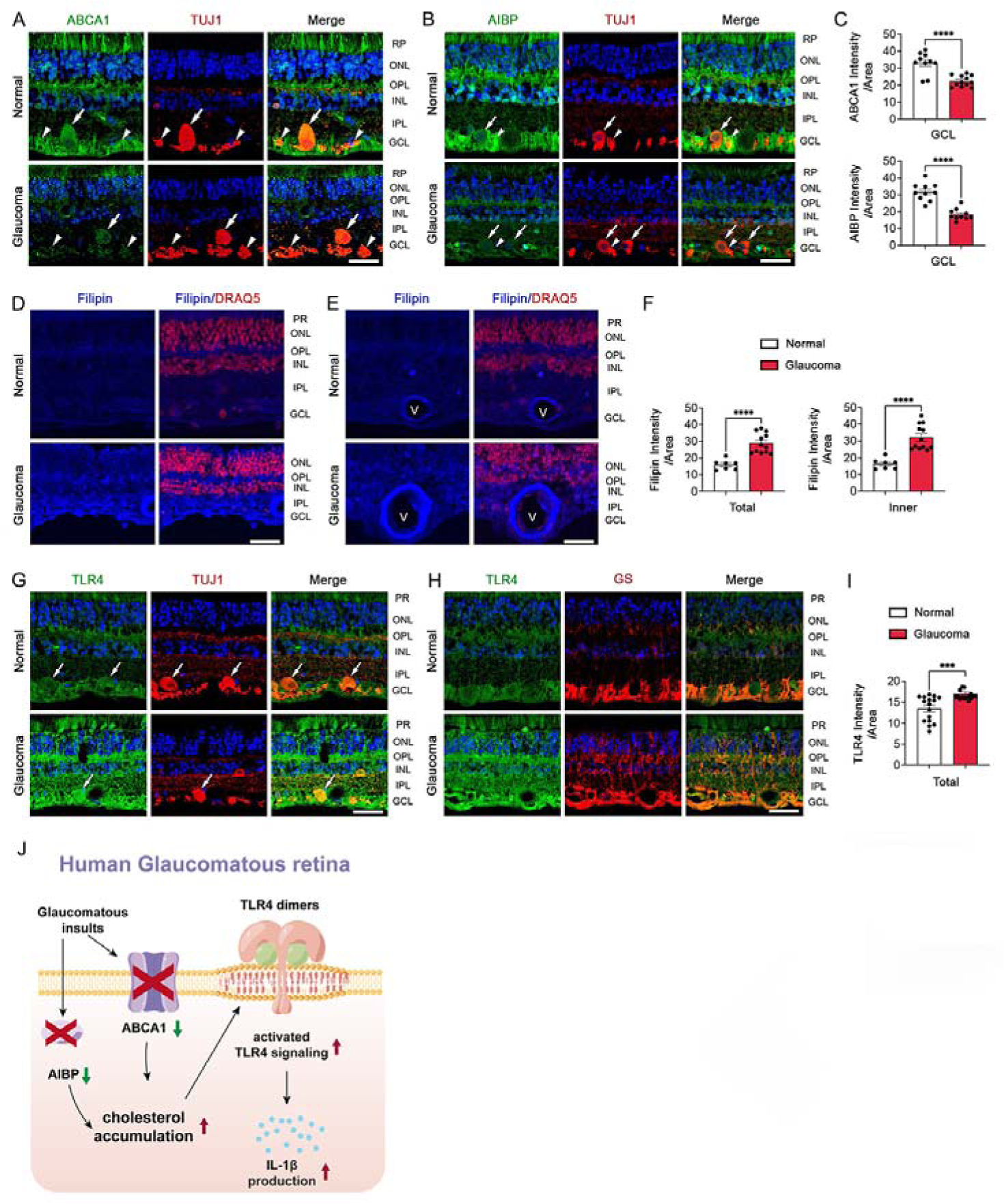
Glaucomatous human retina displays decreased AIBP and ABCA1 protein expression and increased cholesterol content and TLR4 and IL-1β expression. (**A**) Representative retina images for ABCA1 (green) and TUJ1 (red) immunoreactivities. Arrows and arrowheads indicate ABCA1 immunoreactivity co-labeled with TUJ1 in RGC somas and RGC axon bundle, respectively. (**B**) Representative retina images for AIBP (green) and TUJ1 (red) immunoreactivities. Arrows and arrowheads indicate AIBP immunoreactivity co-labeled with TUJ1 in RGC somas and RGC axon bundle, respectively. (**C**) Quantitative fluorescent intensity of ABCA1 and AIBP immunoreactivities in the GCL of the glaucomatous retina. (**D**) Representative retina images for filipin (blue) and DNA (DRAQ5, red) staining for total retinal layer. (**E**) Representative retina images for filipin (blue) and DNA (DRAQ5, red) staining for retinal blood vessels. (**F**) Quantitative fluorescent intensity of filipin staining in the total and inner layers of the glaucomatous retina. (**G**) Representative retina images for TLR4 (green) and TUJ1 (red) immunoreactivities. Arrows indicate TLR4 immunoreactivity co-labeled with TUJ1 in RGC somas. (**H**) Representative retina images for TLR4 (green) and GS (red) immunoreactivities in Müller glia. (**I**) Quantitative fluorescent intensity of TLR4 immunoreactivity in the total layer of the glaucomatous retina. (**J**) Schematic for the effect of increased levels of cholesterol on TLR4/IL-1β activation in the human glaucomatous retina. *n* = 10-16 sections from 2 eyes from control subject, and *n* = 10-16 sections from 4 eyes from patients with glaucoma. Error bars represent SEM. Statistical analysis was performed using Student’s *t*-test. \*\*\**P* < 0.001 and \*\*\*\**P* < 0.0001. Scale bars, 20 μm.

### Increased cholesterol content and TLR4/IL-1**β** expression in human glaucomatous retina

To test whether excessive retinal accumulation of cholesterol occurs in human glaucoma, we performed filipin fluorescent staining. Unesterified cholesterol detected by filipin staining was significantly increased in human glaucomatous compared to normal retina and in the retinal vasculature, suggesting dysregulation of retinal cholesterol metabolism in human glaucoma (Figs. 1 D-F). Cholesterol accumulation was associated with increased TLR4 and IL-1β expression in human glaucomatous retina, with a specific pattern of expression in RGC soma and axons, and in Müller glia (Figs. 1 G-H; *SI Appendix, Figs. S2 C and D*). These results suggest that downregulation of ABCA1 and AIBP expression in glaucoma results in increased levels of retinal cholesterol, which in turn leads to plasma membrane reorganization to maintain lipid rafts. Enlarged and stable lipid rafts support increased TLR4 activation and IL-1β expression, contributing to retinal neuroinflammation (Fig. 1J).

### Restoring AIBP expression protects RGCs and their axons in glaucomatous mice

Based on our findings of reduced AIBP expression in human glaucomatous retina (Figs. 1 B and C; *SI Appendix,* Fig. S2B) and in a mouse model of glaucoma (13), we tested the hypothesis that restoring AIBP expression via intravitreal AAV-AIBP delivery would reduce RGC death in glaucomatous DBA/2J (further referred to as D2) mice. We intravitreally injected AAV-Null or AAV-AIBP at the age of 5 months and then assessed RGC survival and axon preservation at the age of 10 months (Fig. 2A). We used D2-*Gpnmb^+^* mice, which do not develop a glaucoma-like pathology, injected with AAV-Null, as an age-matched control. To reduce the effect of IOP variation in this model, we further selected mice that have had IOP in the range of 20-35 mmHg (Fig. 2B; *SI Appendix, Table S1*) (8). We found that AIBP expression was restored in AIBP-D2 mice but not Null-D2 mice (Figs. 2 C and D).

**Fig. 2.**
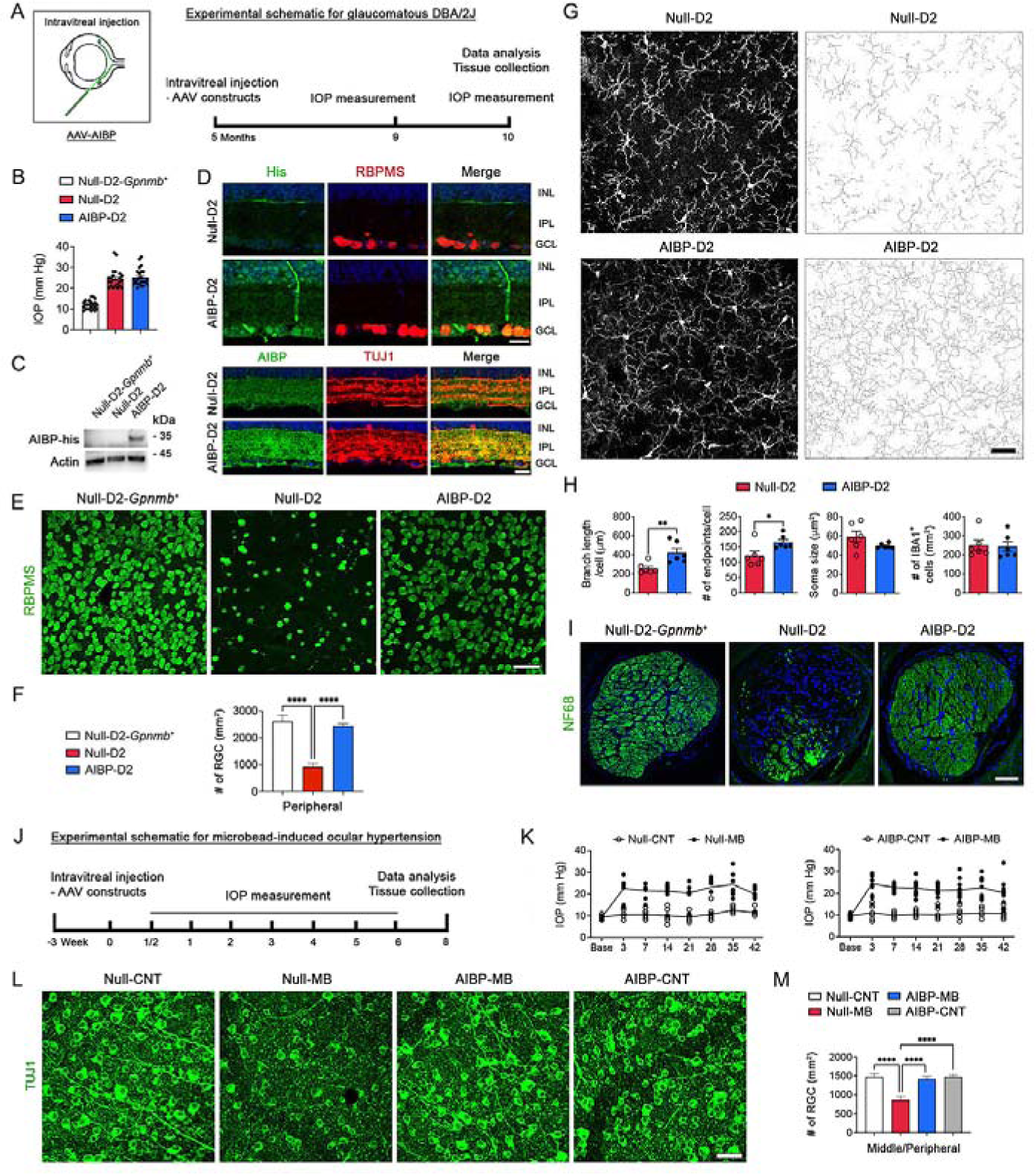
AIBP expression protects RGCs in mouse models of glaucoma. (**A**) Experimental schematic and timeline of AAV injection, IOP measurement, tissue collection and data analysis in D2 mice. (**B**) IOP measurement in 10-month-old D2-*Gpnmb^+^*, Null-D2 and AIBP-D2 mice (*n* = 24 mice per group). (**C**) Representative image of western blot for AIBP expression, detected with an anti-His antibody, in the retinas of Null-D2-*Gpnmb^+^*, Null-D2 and AIBP-D2 mice (*n* = 3 retinas from 3 mice per group). (**D**) Representative retinal images for His (green) and RBPMS (red) immunoreactivities or AIBP (green) and TUJ1 (red) immunoreactivities in Null-D2 and AIBP-D2 mice. (**E**) Representative retina wholemount images for RBPMS (green)-positive RGCs in the peripheral retina. (**F**) Quantitative analysis of RGC numbers in the peripheral areas of the retina (*n* = 5 retina wholemounts from 5 mice for age-matched control Null-D2-*Gpnmb^+^* mice, *n* = 8 retina wholemounts from 8 mice for glaucomatous Null-D2 mice and *n* = 12 retina wholemounts from 12 mice for glaucomatous AIBP-D2 mice). (**G**) Representative retinal wholemount images for IBA1-positive microglial cells in the middle retina of Null-D2 and AIBP-D2 mice. (**H**) Quantitative analysis of branch length (per cell), number of endpoints (per cell), soma size and IBA1-positive number in microglial cells the middle retina (*n* = 6 retinal wholemounts from 6 mice for Null-D2 and AIBP-D2 mice). (**I**) Representative ONH images for NF68 (green)-positive axons in Null-D2-*Gpnmb^+^*, Null-D2, and AIBP-D2 mice. (**J**) Experimental schematic and timeline of AAV transduction, IOP measurement, tissue collection and data analysis in a mouse model of microbead-induced ocular hypertension. (**K**) Monitoring of IOP elevation in MB-injected C57BL/6J mice (*n* = 9-11 mice per group). (**L**) Representative whole-mounted retina images for TUJ1 (green) immunoreactivity in the middle/peripheral area from MB-injected C57BL/6J mice. (**M**) Quantitative analysis of RGC number in the middle/peripheral areas of the retina (*n* = 5 retina wholemounts from 5 mice for age-matched Null-CNT and AIBP-CNT mice; *n* = 7 retina wholemounts from 7 mice for Null-MB mice; *n* = 9 retinal wholemounts from 9 mice for AIBP-MB mice). Error bars represent SEM. Statistical analysis was performed using one-way ANOVA and Tukey’s multiple comparisons test. \*\**P* < 0.01 and \*\*\*\**P* < 0.0001. Scale bars, 20 μm (D) and 50 μm (E and G).

In comparison with non-glaucomatous Null-D2-*Gpnmb^+^* mice, elevated IOP resulted in a significant loss of RGCs in glaucomatous Null-D2 mice (Figs. 2 E and F; *SI Appendix, Table S1*). Remarkably, administration of AAV-AIBP significantly reduced RGC death in D2 mice compared with Null-D2 mice (Figs. 2 E and F; *SI Appendix, Table S1*). We further found that administration of AAV-AIBP significantly increased microglial branch length and number of endpoints in the middle area of the retina, without changes in the soma size or the number of IBA1-positive microglia (Figs. 2 G and H), suggesting that AIBP protects RGCs by preventing microglia activation in glaucomatous neurodegeneration. Furthermore, administration of AAV-AIBP also preserved RGC axons in the ONH (glial laminar area) in D2 mice (Fig. 2I).

We next examined effect of restoring AIBP expression on RGC protection in a mouse model of microbead (MB)-induced ocular hypertension (26, 27). We intravitreally injected six-week-old C57BL/6J mice with AAV-Null or AAV-AIBP at 3 weeks before the MB injection. We then measured IOP weekly and counted RGC numbers at 8 weeks after MB injection (Figs. 2 J and K; *SI Appendix, Table S2*). In comparison with non-glaucomatous AAV-Null-control (Null-CNT) mice, MB-induced IOP elevation in AAV-Null mice (Null-MB) resulted in a significant loss of RGCs (Figs. 2 L and M; *SI Appendix, Table S2*). Importantly, administration of AAV-AIBP significantly reduced RGC death in MB mice (Figs. 2 L and M; *SI Appendix, Table S2*).

### Restoring AIBP expression ameliorates visual dysfunction in glaucomatous axon injury

Optic nerve crush (ONC) is often used as a model of traumatic optic neuropathy or glaucomatous axon injury because it shows similar pathological phenotypes of RGC soma and axon degeneration following the injury. To examine the effect of restoring AIBP expression on RGC protection in ONC injury, we intravitreally injected AAV-Null or AAV-AIBP into the eyes of 4-month-old C57BL/6J mice at 3 weeks before ONC and then counted RGC numbers at 1 week after ONC (Fig. 3A). Consistent with other experimental models of glaucoma, administration of AAV-AIBP significantly reduced RGC death in ONC mice (Figs. 3 B and C; *SI Appendix, Table S3*). To determine the effect of restoring AIBP expression on visual function, we next measured 1) RGC function using the pattern electroretinogram (PERG), a measurement of the electrical activity by RGCs during exposure to a visual stimulus and 2) the maximum spatial frequency that could elicit head tracking (“acuity”) in a virtual-reality optomotor system. In comparison with AAV-Null-CNT mice, ONC resulted in a significant loss of PERG amplitudes in Null-ONC mice (Figs. 3 D and E). Importantly, administration of AAV-AIBP significantly recovered PERG amplitudes in AIBP-ONC mice (Figs. 3 D and E). In addition, ONC resulted in a significant reduction of visual acuity by decreasing spatial frequency in Null-ONC (Fig. 3 F). However, administration of AAV-AIBP significantly restored visual acuity by increasing spatial frequency in AIBP-ONC mice (Fig. 3 F).

**Fig. 3.**
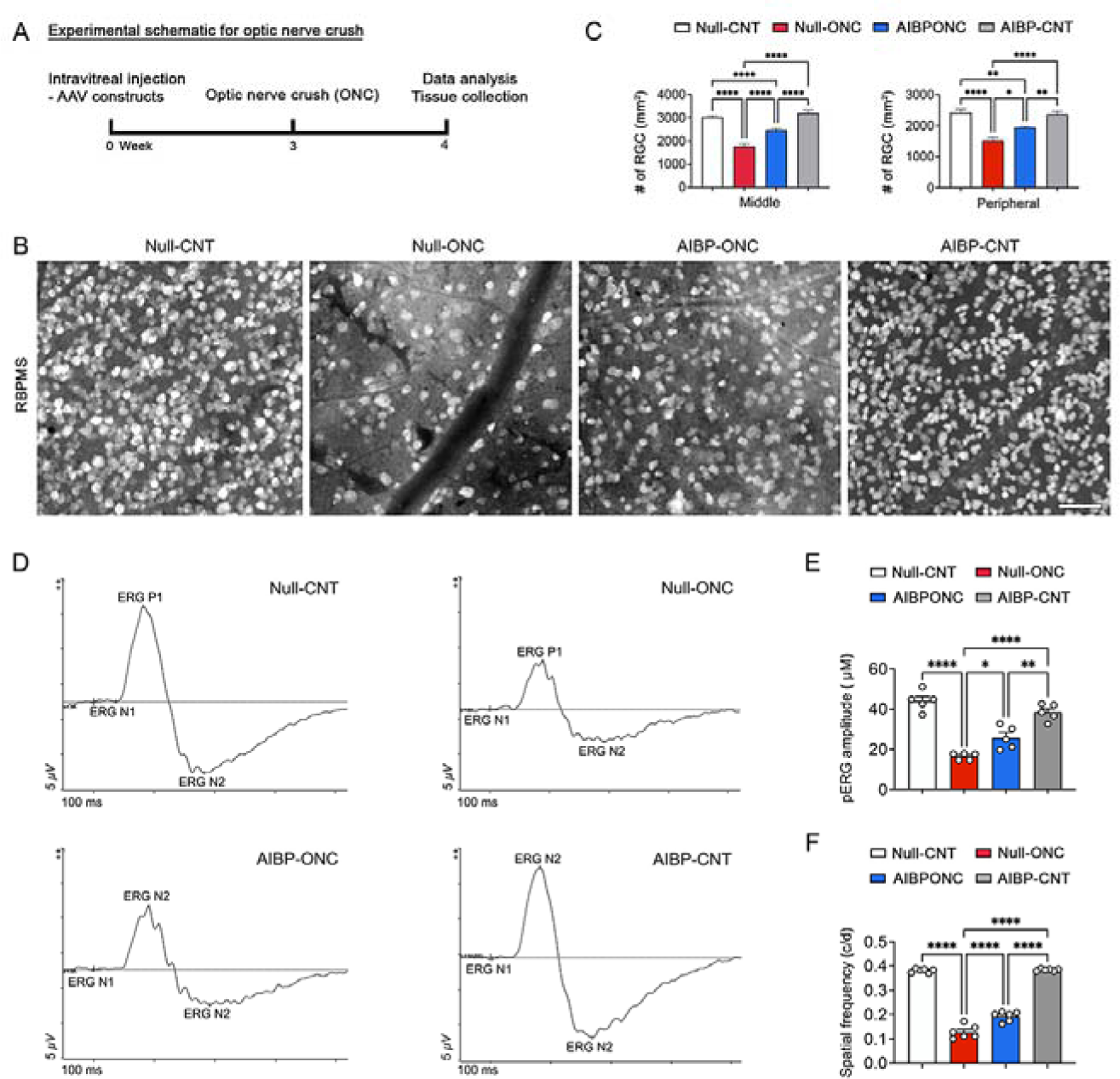
AIBP expression protects RGCs in a mouse model of optic nerve crush. (**A**) Experimental schematic and timeline of AAV transduction, ONC, tissue collection and data analysis in the mouse model of ONC. (**B**) Representative whole-mounted retinal images for RBPMS (gray) immunoreactivities in the middle and peripheral area from ONC-induced C57BL/6J mice. (**C**) Quantitative analysis of RGC number in the middle/peripheral areas of the retina (*n* = 5 retina wholemounts from 5 mice per group). (**D**) Representative recording graphs of pERG amplitude. (**E**) pERG amplitudes analyses (*n* = 5 mice per group). (**F**) Spatial frequency by optomotor response analyses (*n* = 5 mice per group). Note that AIBP expression restores visual function in ONC mice. Error bars represent SEM. Statistical analysis was performed using one-way ANOVA and Tukey’s multiple comparisons test. \**P* < 0.05, \*\**P* < 0.01, \*\*\**P* < 0.001 and \*\*\*\**P* < 0.0001. Scale bar, 50 μm.

### AIBP reduces cholesterol deposition and inhibits TLR4/IL-1**β** expression in the retina of glaucomatous D2 mice

We found decreased TUJ1 immunoreactivity in RGC neurites in the inner plexiform layer and in the RGC somas and axons of the GCL of Null-D2 mice that was significantly increased by administration of AAV-AIBP (Figs. 4 A and B). Since AIBP can facilitate removal of excessive cholesterol from activated cells (20, 28, 29) and because we found an increased cholesterol deposition in the retina of unchallenged AIBP-deficient mice (*SI Appendix,* Fig. S3), we next examined whether administration of AAV-AIBP reduces excessive cholesterol accumulation in the glaucomatous D2 retina. We observed a significant increase of cholesterol deposition in all retinal layers of glaucomatous Null-D2 mice compared with Null-D2-*Gpnmb^+^* mice, which was significantly decreased in AIBP-D2 mice (Figs. 4 C and D), suggesting that restoring retinal AIBP expression can reduce cholesterol deposition and protect RGC somas and dendrites, even under conditions of elevated IOP.

**Fig. 4.**
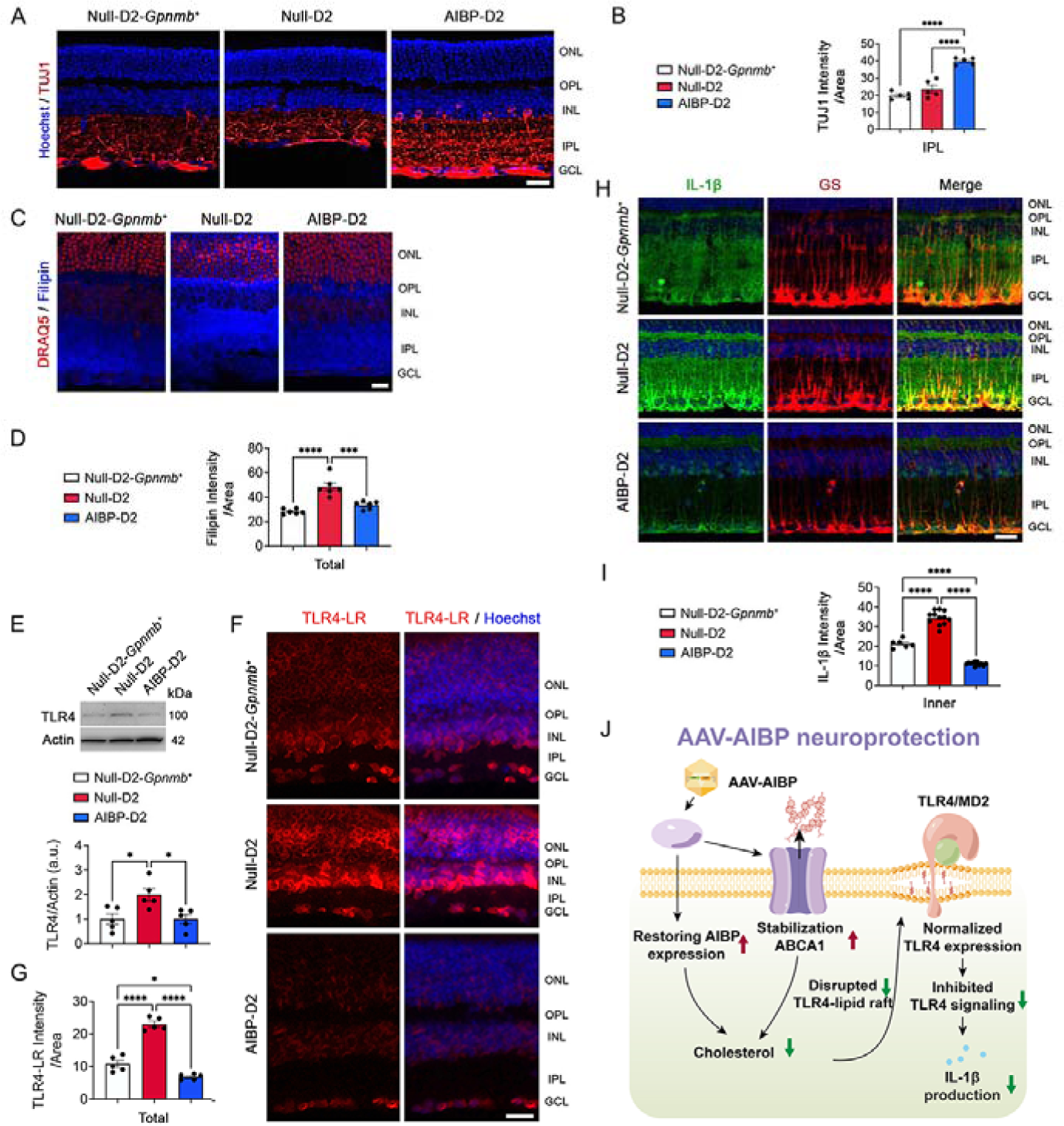
AIBP expression decreases cholesterol deposition and TLR4 and IL-1β expression in glaucomatous D2 retina. (**A**) Representative retina images for TUJ1 immunoreactivity. (**B**) Quantitative fluorescent intensity of TUJ1 intensity in the IPL of the glaucomatous retina (*n* = 5 retina sections from 3 mice per group). (C) Representative retina images for filipin (blue) and DNA (DRAQ5, red) staining for total retinal layer. (**D**) Quantitative fluorescent intensity of filipin staining in the total and inner layers of the glaucomatous retina (*n* = 6 retina sections from 3 mice per group). (**E**) Western blot analysis for TLR4 protein expression in the retinas of 10-month-old D2 mice (*n* = 5 mice per group). (**F**) Representative retina images of PLA detection of TLR4-lipid raft (LR) proximity (red). (**G**) Quantitative fluorescent intensity of PLA detection of TLR4-LR proximity in the total layer of the glaucomatous retina (*n* = 5 retina sections from 3 mice per group). (**H**) Representative retina images for IL-1β (green) and GS (red) immunoreactivities in Müller glia. (**I**) Quantitative fluorescent intensity of IL-1β immunoreactivity in the total layer of the glaucomatous retina (*n* = 6 retina sections from 3 mice per group). (**J**). Schematic for RGC survival by AAV-AIBP-induced reduction of retinal cholesterol deposition and of TLR4-LR formation. Error bars represent SEM. Statistical analysis was performed using one-way ANOVA and Tukey’s multiple comparisons test. \**P* < 0.05, \*\*\**P* < 0.001 and \*\*\*\**P* < 0.0001. Scale bars, 20 μm.

Because glaucomatous D2 retina shows significant increase in TLR4 expression (13) and AIBP plays a role in disrupting TLR4 signaling via removal of excessive cholesterol from activated cells in neuropathic pain states (14, 22), we further examined whether restoring AIBP expression inhibits activation of TLR4 in glaucomatous D2 retina. We observed a significant increase of TLR4 expression, TLR4-associated lipid rafts as measured by a proximity ligation assay, and IL-1β expression in glaucomatous Null-D2 retina compared with Null-D2-*Gpnmb^+^* retina, which were all significantly reduced in AIBP-D2 retina (Figs. 4 E-I). These findings suggest that restoring retinal AIBP expression reduces cholesterol deposition and TLR4-lipid raft formation and inhibits TLR4/IL-1β expression, leading to inhibition of neuroinflammation and RGC protection in experimental mouse models of glaucoma (Fig. 4J).

### AIBP expression reduces MAPK and AMPK activation in the retina of glaucomatous D2 mice

Because MAP kinase-mediated signaling pathways are associated with the activation of retinal glial cells (astrocytes or Müller glia) in the glaucomatous human retina (30), we first examined the expression levels and distribution of phospho-p38 (pp38) and phospho-ERK1/2 (pEKR1/2) in the glaucomatous human retina. In the control human retina, we observed that pp38 immunoreactivity was present in Müller glia and neuronal cells in the inner nuclear layer (INL) (*SI Appendix,* Fig. S4A and B). In the glaucomatous human retina, we found an increase of pp38 immunoreactivity in RGCs and Müller glia (*SI Appendix,* Fig. S4A-C). Consistently, we found a significantly increased pp38 in Null-D2 retina compared with Null-D2-*Gpnmb^+^* retina in RGCs (*SI Appendix,* Fig. S4D-F). Administration of AAV-AIBP significantly reduced pp38 in the inner retinal layer and TUJ1-positive RGCs in AIBP-D2 retina (*SI Appendix,* Fig. S4D-F). In addition, in the control human retina, we observed that pERK1/2 immunoreactivity was present in Müller glia (*SI Appendix,* Fig. S4H), and that it was increased in Müller glia in the inner retinal layers of the glaucomatous human retina (*SI Appendix,* Fig. S4H and I). Interestingly, we did not detect pERK1/2 immunoreactivity in RGCs from both normal and glaucomatous human retinas (*SI Appendix,* Fig. S4G). Consistent with human data, we found significantly increased pERK1/2 in Null-D2 retina compared with Null-D2-*Gpnmb^+^* retina (*SI Appendix,* Fig. S4J), which was reduced to normal levels in AIBP-D2 retina (*SI Appendix,* Fig. S4J). As in human retina, pERK1/2 was detected in other than TUJ1-positive cells (*SI Appendix,* Fig. S4J). These results were replicated in pERK1/2 IHC analyses (*SI Appendix,* Fig. S4K and L).

AMPK is a highly conserved energy sensor and its hyperactivation is prominent in RGCs from experimental glaucoma and patients with primary open-angle glaucoma (POAG) (31). Attenuation of AMPK hyperactivation promotes RGC function and survival by inhibiting dendritic retraction and synaptic loss in magnetic microbead occlusion mouse model of glaucoma (32). We observed that pAMPK (Thr172) immunoreactivity in Müller glia and RGCs was significantly increased in human glaucomatous compared to normal retina (Figs. 5 A-C). Consistent with the human data, we found significantly increased pAMPK in Null-D2 retina compared with Null-D2-*Gpnmb^+^*retina (Fig. 5D). Restoring AIBP expression significantly decreased pAMPK in glaucomatous AIBP-D2 retina as measured by western blot (Fig. 5D). IHC analysis also showed increased pAMPK in Null-D2 RGCs compared with Null-D2-*Gpnmb^+^* RGCs (Figs. 5 E and F) and that restoring AIBP expression decreased pAMPK in both OPL and inner retinal layers of AIBP-D2 mice (Figs. 5 E and F). These results support the notion that retinal AIBP expression delivered by an AAV protects RGCs and improves axonal transport via inhibition of inflammation and restoration of energy homeostasis (Fig. 5G).

**Fig. 5.**
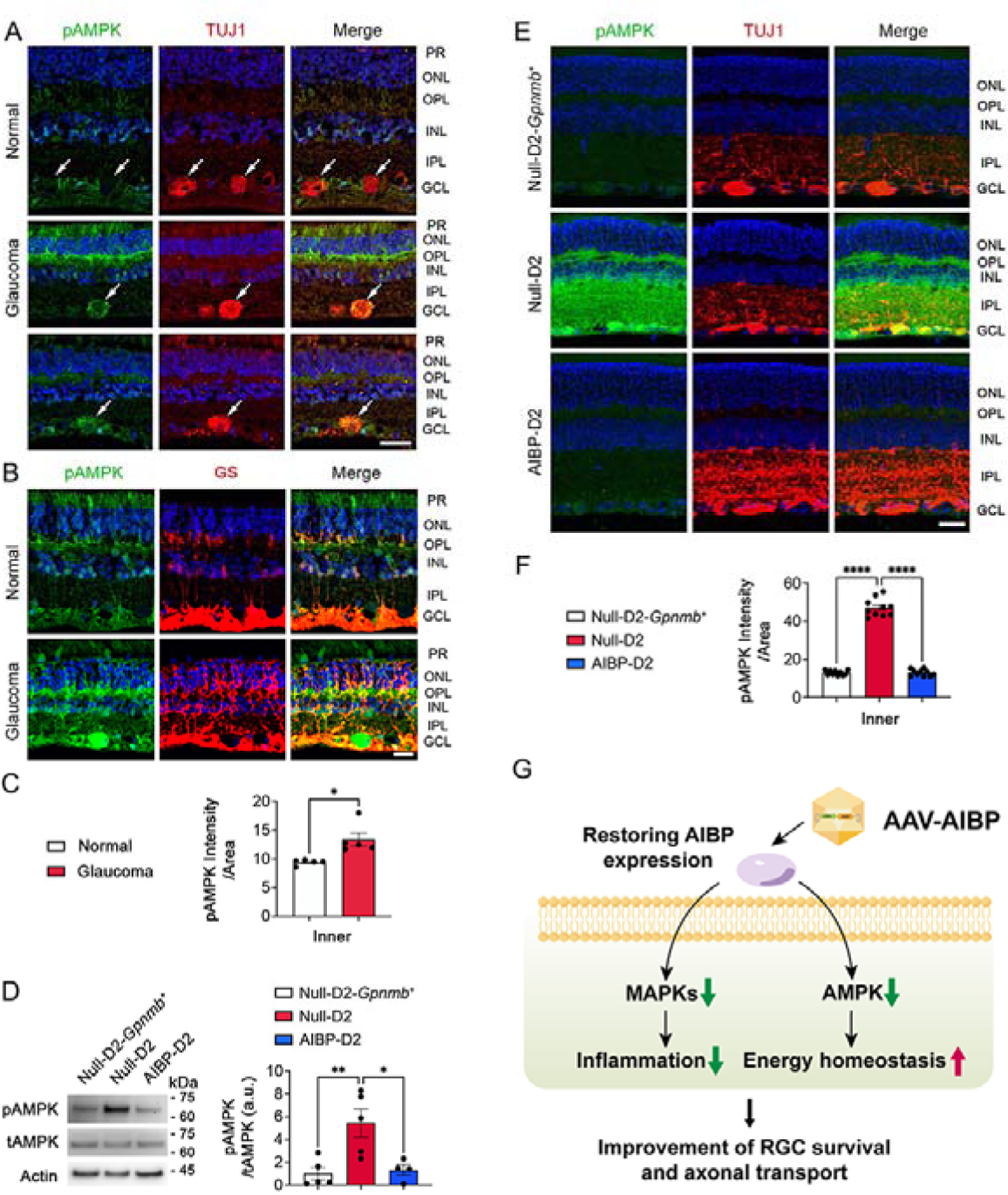
AIBP expression inhibits AMPK hyperactivation in the glaucomatous retina. (**A**) Representative retina images for phospho-AMPK (pAMPK, green) and TUJ1 (red) immunoreactivities in the glaucomatous human retina. (**B**) Representative retina images for pAMPK (green) and GS (red) immunoreactivities in Müller glia of the glaucomatous human retina. (**C**) Quantitative fluorescent intensity of pAMPK immunoreactivity in the inner layer of the glaucomatous human retina (*n* = 5 retina sections from 2 eyes from control subject and *n* = 5 retina sections from 4 eyes from patients with glaucoma). (**D**) Western blot analysis of pAMPK in glaucomatous D2 retina (*n* = 4-5 retinas from 4-5 mice per group). (**E**) Representative retina images for pAMPK (green) and TUJ1 (red) immunoreactivities in glaucomatous D2 retina. (**F**) Quantitative fluorescent intensity of pAMPK immunoreactivity in the inner retina of glaucomatous D2 mice (*n* = 10 retina sections from 3 mice per group). (**G**) Schematic for reduction of retinal MAPKs and AMPK activations by AAV-AIBP. Error bars represent SEM. Statistical analysis was performed using Student’s *t*-test or one-way ANOVA and Tukey’s multiple comparisons test. \**P* < 0.05, \*\**P* < 0.01, and \*\*\*\**P* < 0.0001. Scale bars, 20 μm.

## Discussion

In this study, we report novel findings of significantly reduced ABCA1 and AIBP expression as well as increased cholesterol accumulation that are associated with increasing of TLR4 and IL-1β expression in the retina of human patients with glaucoma. These findings were replicated in a mouse experimental model of glaucoma. The excessive retinal cholesterol deposition was reversed by restoring AIBP expression via a single intravitreal injection of AAV-AIBP. This treatment reduced TLR4 and IL-1β expression and it prevented RGC loss and visual dysfunction in mouse models of glaucoma. The neuroprotective AIBP effect was mediated by reducing phosphorylation of AMPK, p38, and ERK1/2, as well as reducing inflammatory response by glaucomatous insults such as elevated pressure.

We identified AIBP as a master regulator of retinal cholesterol content, which in turn regulates retinal neuroinflammation and RGC survival. The causes of reduced AIBP expression in glaucoma are yet to be elucidated, but an AAV-AIBP intervention to restore AIBP expression normalized retinal cholesterol, disrupted TLR4-lipid raft formation, inhibited TLR4 and IL-1β expression and prevented RGS loss. AIBP is a 32 kDa protein, which depending on phosphorylation by PKA, can be secreted or localized to mitochondria (19, 33). The AAV-AIBP construct employed in our studies contains a strong secretion signal (fibronectin signal peptide), thus the observed effects can be assigned to a secreted form of AIBP. We and others have reported that the secreted, extracellular AIBP facilitates removal of excess cholesterol from endothelial cells and macrophages, resulting in reduction of lipid rafts and inhibition of angiogenesis, atherosclerosis and pulmonary inflammation (20, 21, 28, 29, 34). More recently, we have shown that AIBP binds to TLR4 and mediates selective cholesterol depletion and disruption of lipid rafts in TLR4-expressing cells. The loss of lipid raft integrity in microglia results in inhibition of TLR4 dimerization and reduced spinal neuroinflammation (22, 35). Moreover, AIBP downregulates the surface expression and function of TLR4 and downstream inflammatory pathways in macrophages and microglia (14, 36, 37). In this study, we observed AIBP-mediated reductions in retinal TLR4 expression and TLR4 localization to lipid rafts.

It is established that TLR4 activation simultaneously triggers the activation of MAPK/NF-κB and inflammasome pathways, thereby escalating the secretion of proinflammatory cytokines (4, 5, 38). In the current study, we found that phosphorylation of p38 was increased in both RGCs and Müller glia in the glaucomatous human retina, while phosphorylation of ERK1/2 was increased only in Müller glia. The MAPK signaling has been known to regulate not only inflammatory response (39) but also mitochondrial biogenesis (40, 41). Here, we found that AIBP administration significantly reduced phosphorylation of p38 and ERK1/2 in the inner retinal layer of the glaucomatous D2 mice. Together, these findings suggest that the AIBP administration leading to normalization of TLR4-associated lipid rafts may preserve mitochondrial function and inhibit inflammatory responses by preventing activation of the MAPK pathways under glaucomatous injury and thus protecting RGCs.

Human glaucomatous retina and the retinas in mouse models of glaucoma are characterized by biomarkers of energy depletion and metabolic vulnerability (32, 42, 43), which are associated with inflammatory response (43). AMPK is a highly conserved metabolic energy sensor activated by a variety of energy stresses and mitochondrial damage that reduce ATP levels (44, 45). Because mitochondrial dysfunction produces a low level of ATP and leads to axonal and dendritic degeneration in RGCs (13, 32), AMPK activation would be a pathophysiological indicator for glaucomatous RGC degeneration. Indeed, AMPK activation has been found in experimental mouse models of glaucoma (32, 43). Here, we showed that AIBP administration significantly inhibited AMPK activation in RGCs and Müller glia in the glaucomatous mouse retina, implying that AIBP may protect RGCs and Müller glia by ameliorating energy crisis and metabolic stress induced by glaucomatous insults.

In conclusion, we propose the novel paradigm that controlling TLR4-lipid rafts in the retina – via maintenance of retinal expression of secreted AIBP – can protect RGCs under the conditions of elevated IOP. The mechanism includes inhibition of TLR4-lipid raft activation, and reduced activation of glaucomatous Müller glia, leading to RGC protection and preservation of visual function *(SI Appendix, Figs. S5*).

## Materials and Methods

### Human tissue samples

Human retina tissue sections were obtained, via San Diego Eye Bank, from two eyes of a donor (age 85 years) without history of glaucoma or any other eye disease, and of glaucoma patients (ages 76 and 84 years). The studies were approved by the University of California, San Diego Human Research Protection Program. Both control and glaucoma patients had no history of other eye disease, diabetes, or chronic central nervous system disease.

### Animals

Adult male and female DBA/2J and age-matched D2-*Gpnmb^+^*mice (The Jackson Laboratory), and WT and *Apoa1bp^-/-^* mice were housed in covered cages, fed with a standard rodent diet ad libitum, and kept on a 12 h light/12 h dark cycle. C57BL/6J mice were initially purchased from the Jackson Laboratory, bred in-house for experiments and used as WT mice. *Apoa1bp^-/-^* mice on a C57BL/6J background were generated in our group as previously reported (28, 34). Animals were assigned randomly to experimental and control groups. All procedures concerning animals were in accordance with the Association for Research in Vision and Ophthalmology Statement for the Use of Animals in Ophthalmic Vision Research and under protocols approved by Institutional Animal Care and Use Committee at the University of California, San Diego and the University of Texas Medical Branch.

### Production of recombinant AAV-AIBP vector

AAV-AIBP was generated as previously reported (46). Murine AIBP (25-283aa) was fused with FIB at the N-terminus and 6x-His at the C-terminus (FIB-AIBP-His). FIB-AIBP-His was transduced by the AAV-DJ/8 Helper-free system (Cell BioLabs) which is mimicking serotypes AAV8 and AAV9 with the inverted terminal repeats (ITRs) from AAV2 (Agilent Technologies) that contains the CMV promoter and subsequently purified according to published protocol (47). AAV was titrated using quantitative PCR (qPCR) with primers for the ITR sequence (TaKaRa Bio Inc).

### Transduction with recombinant AAV constructs

The 5-month-old pre-glaucomatous DBA/2J and D2-*Gpnmb^+^* mice were anesthetized with an intraperitoneal (IP) injection of a mixture of ketamine/xylazine and topical 1% proparacaine eye drops. A 32-gauge needle was used to inject a total of 1 µl of either AAV-Null or AAV-AIBP (1×10^13^ gc/ml) into the vitreous humor of the eye. Injections were given slowly over 1 min and the needle was maintained in position for an additional 10 min to minimize vector loss through the injection tract. At the age of 10 months, the mice were euthanized by an IP injection of a mixture of ketamine/xylazine, and the retina and ONH tissues were prepared.

### Mouse model of MB-induced ocular hypertension

MB-induced glaucoma was induced using a modified protocol based on previous publications (26, 27). In brief, six-week-old C57BL/6J mice were intravitreally injected with either AAV-Null or AAV-AIBP as described above. At 3 weeks post-injection, mice were anesthetized by an IP injection of a cocktail of ketamine (100□mg/kg, Ketaset; Fort Dodge Animal Health) and xylazine (9□mg/kg, TranquiVed; VEDCO Inc.). We topically applied 0.5% proparacaine hydrochloride and the pupils were dilated with 1% tropicamide and 2.5% phenylephrine and then the cornea was gently punctured using a 30-gauge needle. Next, 2 µl of 1-µm-diameter polystyrene MB suspension (containing 3.0 × 10^7^ beads) followed by an air bubble, 2 µl of 6-µm-diameter polystyrene MB suspension (containing 6.3 × 10^6^ beads, Polysciences), and 1 µl of PBS containing 30% Healon was injected into the anterior chamber using a 32-gauge needle (Hamilton Company) connected to a syringe (Hamilton Company), which induced moderate elevation of IOP for ≥6 wk. An equivalent volume of PBS was injected into the contralateral eyes to serve as sham control. At 8 weeks after the MB injection, mice were sacrificed, and the retina tissues were prepared.

### IOP measurement

IOP elevation onset in DBA/2J mice typically occurs between 5 and 7 months of age, and by 9 to 10 months of age, IOP-linked optic nerve axon loss is well advanced (8, 9). IOP measurements were performed as previously described (8, 9). Each of the 10-month-old D2-*Gpnmb^+^* and DBA/2J mice used in this study had a single IOP measurement (to confirm development of spontaneous IOP elevation exceeding 20 mmHg) using a tonometer (TonoLab, Colonial Medical Supply) (*n* = 24 mice per group). For the mouse model of MB-induced glaucoma, IOP was measured using a tonometer (TonoLab) from 3 days after MB injection thereafter until 6 weeks after injection between 3 p.m. and 5 p.m. to minimize diurnal variability. Non-IOP elevation contralateral control retinas were used as sham control (*n* = 9-11 mice per group).

### Tissue preparation

Mice were anesthetized by an IP injection of a cocktail of ketamine/xylazine as described above prior to cervical dislocation. For immunohistochemistry, the retinas and ONHs were dissected from the eyeballs and fixed with 4% paraformaldehyde (Sigma) in PBS (pH 7.4) for 2□h at 4□°C. Retinas were washed several times with PBS then dehydrated through graded levels of ethanol and embedded in polyester wax. For EM, the eyes were fixed via cardiac perfusion with 2% paraformaldehyde, 2.5% glutaraldehyde (Ted Pella) in 0.15□M sodium cacodylate (pH 7.4, Sigma) solution at 37□°C and placed in pre-cooled fixative of the same composition on ice for 1□h. As described below, the procedure was used to optimize mitochondria structural preservation and membrane contrast. For western blot analyses, extracted retinas were immediately used.

### Western blot analyses

Harvested retinas were homogenized for 1 min on ice with a RIPA lysis buffer (Cell Signaling Technology), containing protease inhibitor cocktail (MedChemExpress). The lysates were then centrifuged at 13,000 rpm for 15 min and protein amounts in the supernatants were measured by DC Protein assay (Bio-Rad). Proteins (10-20 μg) were run on a NuPAGE Bis-Tris gel (Invitrogen) and transferred to polyvinylidene difluoride membranes (PVDF) membranes (GE Healthcare Bio-Science). The membranes were blocked with TBS/0.1% Tween-20 (T-TBS) containing 5% bovine serum albumin (BSA) for 1□h at room temperature and incubated with primary antibodies for overnight at 4□°C. Membrane were washed three times with T-TBS then incubated with horseradish peroxidase (HRP)-conjugated secondary antibodies (Cell Signaling Technology) for 1 h at room temperature. Membranes were developed using enhanced chemiluminescence substrate system (ThermoFisher Scientific). The images were captured using a UVP imaging system.

### Immunohistochemistry

Immunohistochemical staining of 7 μm wax sections of full thickness retina were performed. Sections from wax blocks from each group were used for immunohistochemical analysis. To prevent non-specific background, tissues were incubated in 1% BSA/PBS for 1 h at room temperature before incubation with the primary antibodies for 16 h at 4°C. After several wash steps, the tissues were incubated with the secondary antibodies for 4 h at 4°C and subsequently washed with PBS. The sections were counterstained with the nucleic acid stain Hoechst 33342 (1 μg/ml; Invitrogen) in PBS. Images were acquired with Keyence All-in-One Fluorescence microscopy (BZ-X810, Keyence Corp. of America). Each target protein fluorescent integrated intensity in pixel per area was measured using the ImageJ software (National Institutes of Health). All imaging parameters remained the same and were corrected with the background subtraction.

### Whole-mount immunohistochemistry and RGC counting

Mouse retinas from enucleated eyes were dissected as flattened whole-mounts. Retinas were immersed in PBS containing 30% sucrose for 24 h at 4°C. The retinas were blocked in PBS containing 3% donkey serum, 1% BSA, 1% fish gelatin and 0.1% triton X-100, and incubated with primary antibodies for 3 days at 4°C. Primary antibodies used were rabbit polyclonal anti-RBPMS antibody or mouse monoclonal anti-TUJ1 antibody. After several wash steps, the retinas were incubated with Alexa Fluor-488 conjugated donkey anti-rabbit IgG antibody or Alexa Fluor-568 conjugated donkey anti-mouse IgG antibody for 24 h, and subsequently washed with PBS. Images were acquired with Keyence All-in-One Fluorescence microscopy (BZ-X810, Keyence Corp. of America). RBPMS- or TUJ1-positive RGCs were counted in four regions within the middle and/or peripheral retina (three sixths for middle and five sixths for peripheral of the retinal radius) and the scores were averaged using ImageJ.

### Microglia morphology analysis

Whole-mounted mouse retinas were stained with IBA-1 antibody and z-stack images were acquired with Olympus FluoView 1000 confocal microscope (Olympus) with 20x magnification. For microglia morphology analysis, branch length, number of endpoints, soma size, and number of microglia were measured using ImageJ as described (48).

### Filipin staining

Fluorescence probe filipin binds to unesterified cholesterol in biological membranes. Frozen retina sections were immersed in PBS for 10 min. The sections were incubated with 1.5 mg/ml glycine (Sigma-Aldrich) for 10 min and followed by staining with 50 µg/ml filipin (Sigma-Aldrich) for 2 h at room temperature in the dark. The sections were washed 3 times with PBS and counter stained with DRAQ5 (BioLegend) to stain DNA for 10 min. Images were acquired with an Olympus FluoView1000 confocal microscopy (Olympus). The fluorescent integrated intensity in pixel per area was measured using the ImageJ. All imaging parameters remained the same and were corrected with the background subtraction.

### PERG analysis

PERG recordings were performed by Celeris ERG platform equipped with pattern ERG stimulators (Diagnosys LLC) (49). Following anesthesia and eye dilation pattern, ERG stimulators were placed on corneal surface. PERG amplitude was measured between first positive peak (P1) and second negative peak (N2).

### Virtual optomotor response analysis

Spatial visual function was performed on a virtual optomotor system (OptoMotry; CerebralMechanics Inc., AB, Canada) (13). Spatial frequency threshold, a measure of visual acuity, was determined automatically with accompanying OptoMotry software, which uses a step-wise paradigm based upon head-tracking movements at 100% contrast.

### Proximity ligation assay (PLA)

The PLA oligo/probe-conjugated mouse anti-TLR4 (BioLegend) and goat anti-cholera toxin subunit B (CTB, Sigma) antibodies were created using Duolink® PLA probemaker (Sigma-Aldrich) according to the manufacturer’s instructions. To measure TLR4-associated lipid raft (GM1 ganglioside) formation, Duolink® Proximity ligation assay (PLA, Sigma) was performed by following the manufacturer’s instructions. Briefly, frozen retina sections were immersed in H_2_O for 5 min at room temperature. The sections were blocked with one drop of Duolink blocking buffer for 1 h at 37°C. For lipid raft staining, the sections were first incubated with 5 µg/ml of unconjugated CTB (Sigma) for 45 min at 37°C. The oligo/probe-conjugated anti-mouse TLR4 and anti-goat CTB antibodies (1:100) were added to the sections and incubated for additional 2 h at 37°C. The sections were washed twice with Duolink wash buffer, incubated with ligation mixture for 30 min 37°C, followed by incubating with amplification mixture overnight at 37°C. Next day, the sections were washed and incubated with detection buffer for 30 min at 37°C. Images were acquired with Keyence All-in-One Fluorescence microscopy. Each TLR4-CTB PLA fluorescent integrated intensity in pixel per area was measured using the ImageJ. All imaging parameters remained the same and were corrected with the background subtraction.

### Statistical analyses

All results were reported as means ± SEM. For comparison between two groups, statistical analysis was performed using a two-tailed Student’s t-test. For multiple group comparisons, we used one-way analysis of variance (ANOVA) and Tukey’s multiple comparisons test, using GraphPad Prism. A *P* value less than 0.05 was considered statistically significant. \**P* < 0.05, \*\**P* < 0.01, \*\*\**P* < 0.001, \*\*\*\**P* < 0.0001. The value of n per group is indicated within each figure legend.

## Acknowledgments

The authors thank Dr. Shenglin Li for creating illustrative diagrams. This work was supported by the National Institutes of Health grants EY031697 (W.K.J.), AG081004 (W.K.J.), EY034116 (W.K.J./K.Y.K/S.H.C), NS129684 (W.K.J.), EY034376 (W.Z.), EY022694 (W.Z.), EY026629 (W.Z.), HL136275 (Y.I.M.), AG081037 (Y.I.M./W.K.J.), and P30 EY022589 (Core Vision Research).

## Author Contributions

Conceptualization: W.K.J., Y.I.M., S.H.C.; Methodology: W.K.J., Y.H., K.Y.K., T.B., S.C., J.K., S.H.C.; Investigation: W.K.J., Y.H., K.Y.K., T.B., S.C., J.K., S.H.C.; Visualization: W.K.J., Y.H., K.Y.K., S.H.C.; Funding acquisition: W.K.J., Y.I.M.; Project administration: W.K.J., S.H.C.; Supervision: W.K.J., S.H.C.; Writing – original draft: W.K.J., K.Y.K., Y.I.M., S.H.C.; Writing – review & editing: W.K.J., Y.H., K.Y.K., R.N.W., W.Z., Y.I.M., S.H.C.

## Classification

Biological Science, Neuroscience.

