## Supplemental Figures and Tables for "Restoring AIBP expression in the retina provides neuroprotection in glaucoma"

**Affiliations:**

**This PDF file includes:**

Supplementary figures 1 to 5

Supplementary Tables 1 to 4

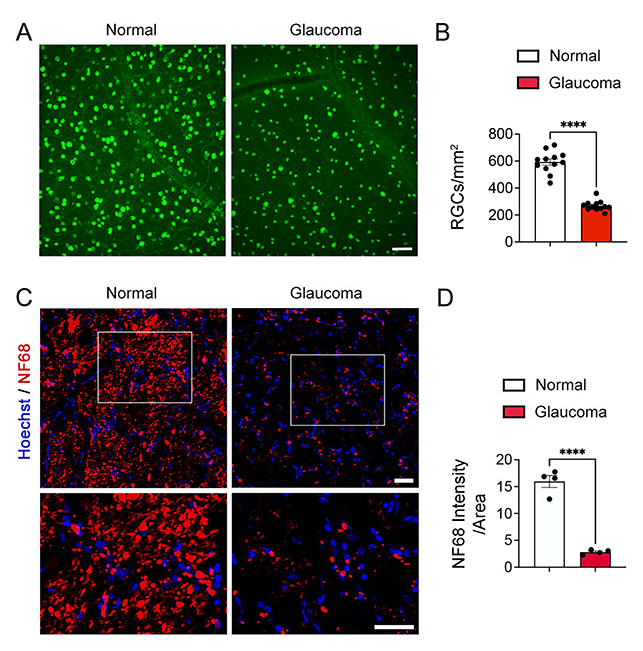

**Fig. S1. Glaucomatous damage induces loss of RGCs and their axons in glaucomatous human retina and ONH** (A) Representative retina wholemount images for RBPMS (green)-positive RGCs in the middle retina. (**B**) Quantitative analysis of RGC numbers in the middle area of the retina (*n* = 12 images from 2 retinas from control subject, and *n* = 12 images from 2 retinas from patients with glaucoma). (**C**) Representative ONH images for NF68 (red) immunoreactivity with nucleus (blue) staining in the laminar cribrosa of the ONHs. (**D**) Quantitative fluorescent intensity showed a significant decrease in NF68 immunoreactivity in the laminar cribrosa of the ONH from patients with POAG (*n* = 4 ONH sectional per group). Error bars represent SEM. Statistical analysis was performed using Student’s *t*-test. *****P* < 0.0001. Scale bars, 100 μm (A) and 20 μm (C).

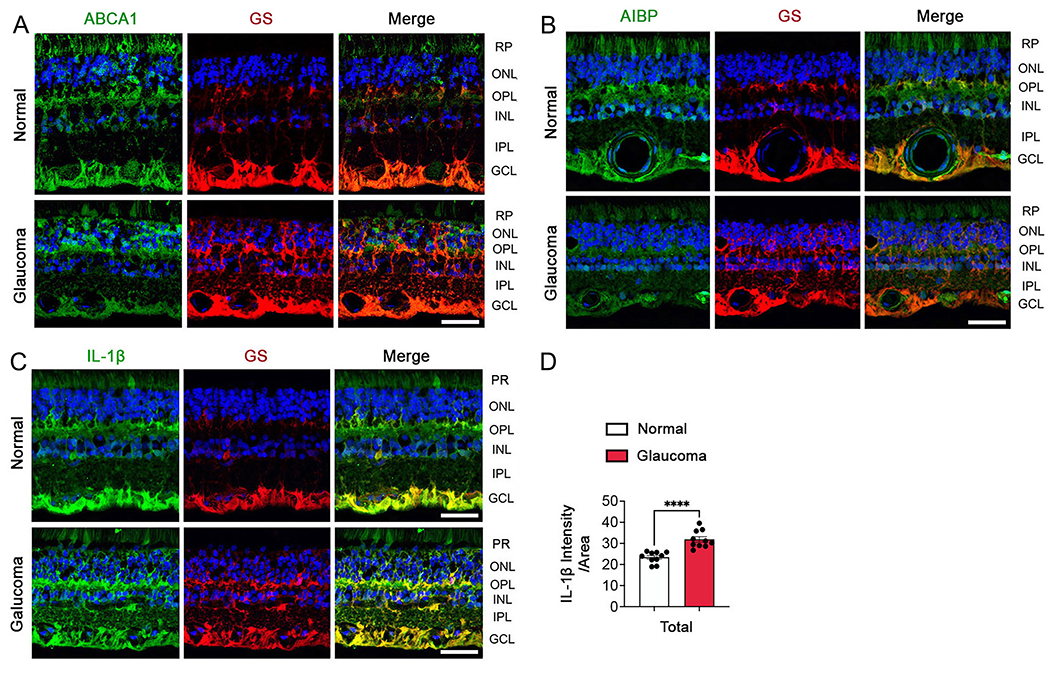

**Fig. S2. Glaucomatous human retina displays increased IL-1β expression and decreased ABCA1 and AIBP protein expression in Müller glia.** (A) Representative retina images for ABCA1 (green) and GS (red) immunoreactivities in Müller glia. (**B**) Representative retina images for AIBP (green) and GS (red) immunoreactivities in Müller glia. (**C**) Representative retina images for IL-1**β** (green) and GS (red) immunoreactivities in Müller glia. (**D**) Quantitative fluorescent intensity of IL-1β immunoreactivity in the total layer of the glaucomatous retina. The sections were counterstained with the nucleic acid stain Hoechst 33342 (blue). *n* = 10-16 sections from 2 eyes from control subject, and *n* = 10-16 sections from 4 eyes from patients with glaucoma. Error bars represent SEM. Statistical analysis was performed using Student’s *t*-test. *****P* < 0.0001. Scale bars, 20 μm.

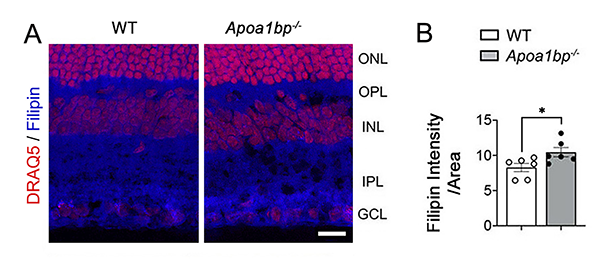

**Fig. S3. AIBP deficiency increases cholesterol content in the retina of *Apoa1bp^-/-^* mice.** (**A**) Representative retina images for filipin (blue) and DNA (DRAQ5, red) staining in the retina from WT and 4-mo-old *Apoa1bp^-/-^* mice. (**B**) Quantitative fluorescent intensity showed a significant increase of filipin intensity in the total retinal layer of Apoa1bp^-/-^ mice (*n* = 6 retina section from 3 mice per group). Error bars represent SEM. Statistical analysis was performed using Student’s *t*-test. **P* < 0.05. Scale bar, 20 μm.

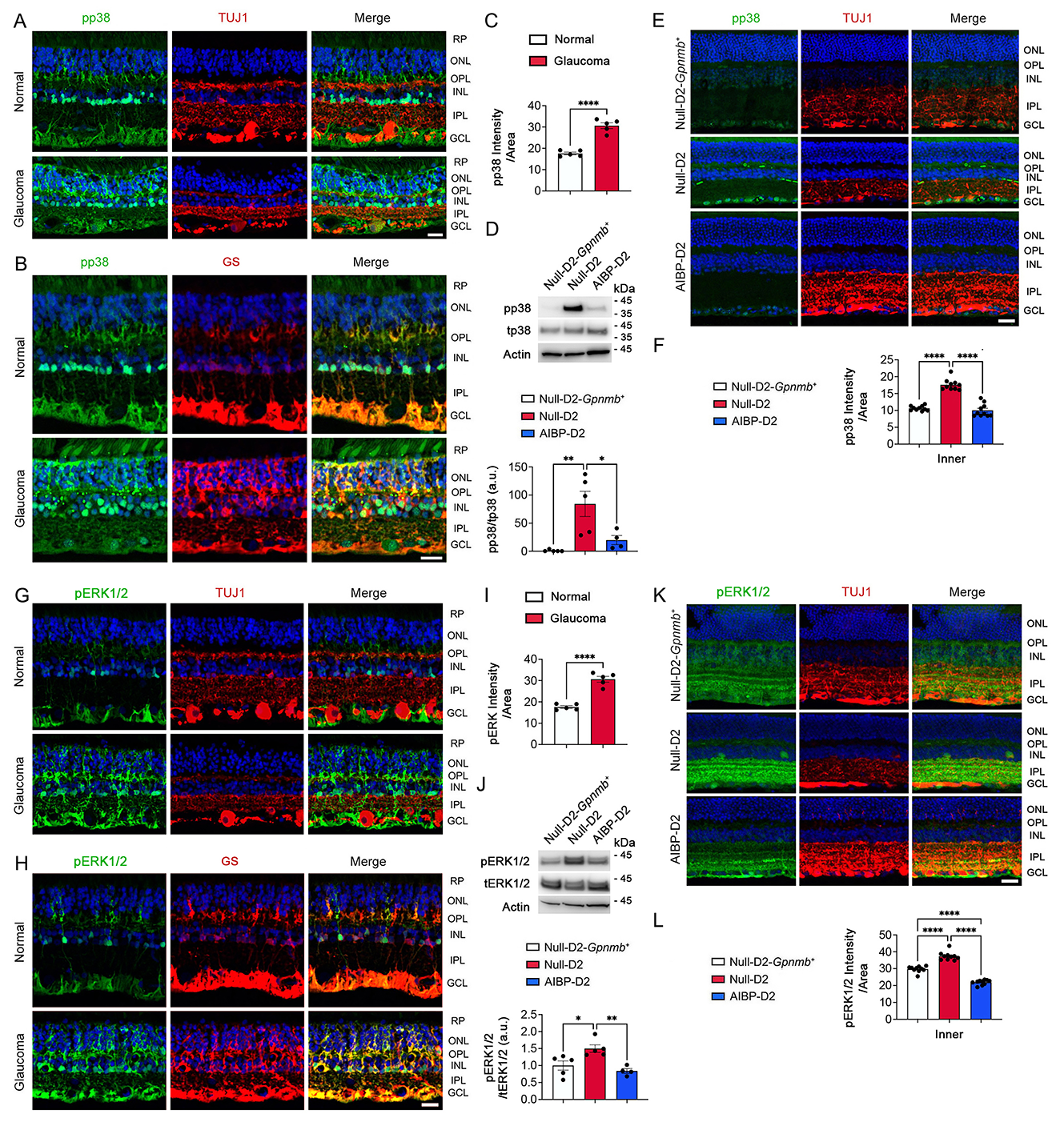

**Fig. S4. AIBP expression inhibits MAPK signaling in the glaucomatous retina.** (**A**) Representative retina images for phospho-p38 (pp38, green) and TUJ1 (red) immunoreactivities in the glaucomatous human retinas. (**B**) Representative retina images for pp38 (green) and GS (red) immunoreactivities in Müller glia of the glaucomatous human retina. (**C**) Quantitative fluorescent intensity of pp38 immunoreactivity in the glaucomatous human retina (*n* = 5 retina sections from 2 eyes from control subject and *n* = 5 retina sections from 4 eyes from patients with glaucoma). (**D**) Western blot analysis of pp38 in glaucomatous D2 retina (*n* = 4-5 retinas from 4-5 mice per group). (**E**) Representative retina images for pp38 (green) and TUJ1 (red) immunoreactivities in glaucomatous D2 retina. (**F**) Quantitative fluorescent intensity of pp38 immunoreactivity in the inner retinal layer of glaucomatous D2 mice (*n* = 10 retina sections from 3 mice per group). (**G**) Representative retina images for phospho-ERK1/2 (pERK1/2, green) and TUJ1 (red) immunoreactivities in the glaucomatous human retinas. (**H**) Representative retina images for pERK1/2 (green) and GS (red) immunoreactivities in Müller glia of the glaucomatous human retina. (**I**) Quantitative fluorescent intensity of pERK1/2 immunoreactivity in the glaucomatous human retina (*n* = 5 retina sections from 2 eyes from control subject and *n* = 5 retina sections from 4 eyes from patients with glaucoma). (**J**) Western blot analysis of pERK1/2 in glaucomatous D2 retina (*n* = 4-5 retinas from 4-5 mice per group). (**K**) Representative retina images for pERK1/2 (green) and TUJ1 (red) immunoreactivities in glaucomatous D2 retina. (**L**) Quantitative fluorescent intensity of pERK1/2 immunoreactivity in the inner retinal layer of glaucomatous D2 mice (*n* = 10 retina sections from 3 mice per group). Error bars represent SEM. Statistical analysis was performed using Student’s *t*-test or one-way ANOVA and Tukey’s multiple comparisons test. **P* < 0.05, ***P* < 0.01, and *****P* < 0.0001. Scale bars, 20 μm.

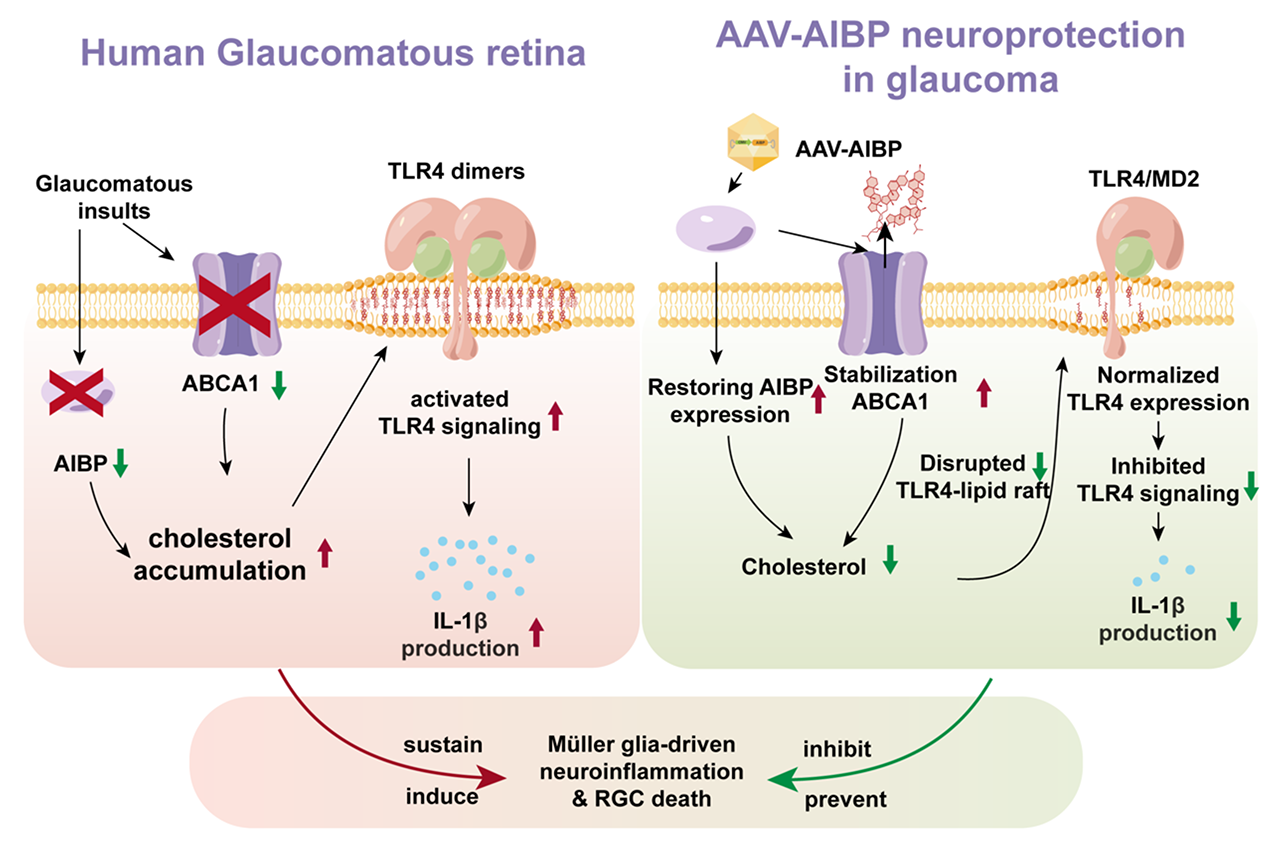

**Fig. S5. AIBP-mediated inhibition of retinal cholesterol accumulation, TLR4-lipid raft activation, inflammatory response, and RGC death in glaucomatous neuroinflammation and neurodegeneration.**

**Table S1. Effect of AAV-AIBP on RBPMS-positive RGC survival in the middle and peripheral retina from 10-month-old glaucomatous DBA/2J mice.**

| Strain | Age  (Months) | Mean IOP (mmHg)  at age of 10 months | RGC density per retina  (RGCs/mm^2^) |
| --- | --- | --- | --- |
|  | |  | Peripheral |
| Null-D2-*Gpnmb^+^* | 10 | 11.8 ± 0.4 | 2608 ± 230 |
| Null-D2 | 10 | 24.5 ± 0.9 | 916 ± 143 |
| AIBP-D2 | 10 | 25.2 ± 0.8 | 2465 ± 92 |

All results were reported as means ± SEM. *n* = 5 retina wholemounts from 5 mice for age-matched control Null-D2-*Gpnmb^+^* mice, *n* = 8 retina wholemounts from 8 mice for glaucomatous Null-D2 mice and *n* = 12 retina wholemounts from 12 mice for glaucomatous AIBP-D2 mice.

**Table S2. Effect of AAV-AIBP on TUJ1-positive RGC survival in the middle/peripheral retina from glaucomatous mice induced by microbead (MB)-induced ocular hypertension.**

| Strain | Age  (weeks) | Mean IOP (mmHg)  at 4 weeks | RGC density per retina  (RGCs/mm^2^) |
| --- | --- | --- | --- |
|  | |  | Middle/Peripheral |
| Null-CNT | 6 | 10.7 ± 1.0 | 1474 ± 77 |
| Null-MB | 6 | 23.2 ± 1.7 | 935 ± 98 |
| AIBP-MB | 6 | 21.5 ± 1.4 | 1412 ± 68 |
| AIBP-CNT | 6 | 10.5 ± 0.6 | 1474 ± 53 |

All results were reported as means ± SEM. *n* = 5 retina wholemounts from 5 mice for age-matched Null-CNT and AIBP-CNT mice; *n* = 7 retina wholemounts from 7 mice for Null-MB mice; *n* = 9 retinal wholemounts from 9 mice for AIBP-MB mice.

**Table S3. Effect of AAV-AIBP on RBPMS-positive RGC survival in the middle and peripheral retina from mouse induced by optic nerve crush injury.**

| Strain | Age (Months) | RGC density per retina  (RGCs/mm^2^) | |
| --- | --- | --- | --- |
|  | | Middle | Peripheral |
| Null-CNT | 4 | 3014 ± 66 | 2424 ± 103 |
| Null-ONC | 4 | 1772 ± 118 | 1536 ± 92 |
| AIBP-ONC | 4 | 2475± 60 | 1950 ± 35 |
| AIBP-CNT | 4 | 3199 ± 126 | 2360 ± 119 |

All results were reported as means ± SEM. *n* = 5 retina wholemounts from 5 mice per group.

**Table S4. Key Resources**

| **REAGENT OR RESOURCES** | **SOURCE** | **IDENTIFIER** |
| --- | --- | --- |
| **Antibodies** |  |  |
| ABCA1 | Novus Biologicals | NB400-105 |
| AIBP | Novus Biologicals | NBP2-30626 |
| AIBP (rabbit polyclonal) | Navia-Pelaez et al., 2022 | N/A |
| AMPK | Cell Signaling Technology | 5831 |
| Phospho-AMPK | Cell Signaling Technology | 2535 |
| β-ACTIN | Cell Signaling Technology | 4967 |
| ERK1/2 | Cell Signaling Technology | 1240 |
| Phospho-ERK1/2 | Cell Signaling Technology | 4370 |
| p38 | Cell Signaling Technology | 8690 |
| Phospho-p38 | Cell Signaling Technology | 4511 |
| GS | Proteintech | 11037-2-1P |
| GS | Chemicon | MAB302 |
| HIS | ThermoFisher Scientific | MA1-21315 |
| HIS | ECM Biosciences | HM0501 |
| IBA1 | Wako | 019-19741 |
| IL-1β | Abcam | ab9722 |
| RBPMS | Novus Biologicals | NBP2-20112 |
| RBPMS | phosphosolutions | 1832-RBPMS |
| TLR4 | Proteintech | 19811-1-AP |
| TLR4 | Genway Biotech | GWB-7F3799 |
| TUJ1 | BioLegend | 801202 |
| Goat anti-rabbit HRP | Cell Signaling Technology | 7074 |
| Goat anti-mouse HRP | Cell Signaling Technology | 7076 |
| Alexa Fluor-488 conjugated donkey anti-mouse IgG antibody | Invitrogen | A-21203 |
| Alexa Fluor-568 conjugated donkey anti-mouse IgG antibody | Invitrogen | A-10037 |
| Alexa Fluor-488 conjugated donkey anti-rabbit IgG antibody | Invitrogen | A-21206 |
| Alexa Fluor-568 conjugated donkey anti-rabbit IgG antibody | Invitrogen | A-10042 |
| Alexa Fluor-488 conjugated donkey anti-guinea pig IgG antibody | Jackson ImmunoResearch | 711-545-152 |
| **Bacterial and Virus Strains** |  |  |
| pAAV-DJ/8-FIB-AIBP [NM_144772.3]-His | Dubrovsky et al. | N/A |
| **Biological Samples** |  |  |
| Human retina tissue samples | San Diego Eye Bank | N/A |
| **Chemicals, commercial kits, and recombinant proteins** |  |  |
| Ketamine | Dechra | 000680 |
| Xylazine | VetOne | 510004 |
| Filipin | Sigma-Aldrich | F4767 |
| DRAQ5 | BioLegend | 424101 |
| AAVpro Titration kit | TaKaRa | 6233 |
| Duolink PLA | Sigma-Aldrich | DUO96030 |
| Alexa Fluor 594-conjugated CTB | ThermoFisher Scientific | C34777 |
| Recombinant AIBP protein | Schneider et al., 2018 | N/A |
| **Experimental models : Strains/Cell lines** |  |  |
| C57BL/6J | Jackson laboratory | Stock No: 000664  RRID:IMSR_JAX:000664 |
| DBA/2J | Jackson laboratory | Stock No: 000671  RRID:IMSR_JAX:000671 |
| DBA/2J-*Gpnmb^+^*/SjJ | Jackson laboratory | Stock No: 007048  RRID:IMSR_JAX:007048 |
| *Apoa1bp^-/-^* mouse | Mao et al., 2017;Schneider et al., 2018 | N/A |
| rMC-1 | Kerafast | ENW001 |
| **Recombinant DNA** |  |  |
| pAAV-MCS | Agilent Technologies | 240071 |
| pAAV-DJ/8 | Cell Biolabs, Inc | VPK-420-DJ-8 |
| pHelper | Cell Biolabs, Inc | 340202 |
| **Software and Algorithms** |  |  |
| ImageJ | NIH, USA | https://imagej.nih.gov/ij/ |
| Adobe Photoshop | USA | https://photoshop.com |
| Prism 9 | GraphPad Inc. | https://www.graphpad.com/scientific-software/prism/ |
